# Overexpression of MUC1 influences the anti-proliferative effect of STAT3-inhibitor Napabucasin in epithelial cancers

**DOI:** 10.1101/2023.06.22.546177

**Authors:** Mukulika Bose, Alexa Sanders, Aashna Handa, Aabha Vora, Cory Brouwer, Pinku Mukherjee

## Abstract

MUC1 is a transmembrane glycoprotein that is overexpressed and aberrantly glycosylated in epithelial cancers. The cytoplasmic tail of MUC1 (MUC1 CT) aids in tumorigenesis by upregulating the expression of multiple oncogenes. Signal transducer and activator of transcription 3 (STAT3) plays a crucial role in several cellular processes and is aberrantly activated in many cancers. In this study, we focus on recent evidence suggesting that STAT3 and MUC1 regulate each other’s expression in cancer cells in an auto-inductive loop and found that their interaction plays a prominent role in mediating epithelial-to-mesenchymal transition (EMT) and drug resistance. The STAT3 inhibitor Napabucasin was in clinical trials but was discontinued due to futility. We found that higher expression of MUC1 increased the sensitivity of cancer cells to Napabucasin. Therefore, high-MUC1 tumors may have a better outcome to Napabucasin therapy. We report how MUC1 regulates STAT3 activity and provide a new perspective on repurposing the STAT3-inhibitor Napabucasin to improve clinical outcome of epithelial cancer treatment.

## Introduction

STAT3 belongs to a family of transcription factors comprising STAT1, STAT2, STAT3, STAT4, STAT5A, STAT5B, and STAT6. In humans, this is encoded by the STAT3 gene. STAT3 is activated by several cytokines and growth factors such as IL-6, IL-9, IL-10, IL-27, TNF-α, MCP-1, EGF, PDGF, G-CSF, and GM-CSF. Active STAT3 plays an important role in tumorigenesis by regulating transcription of genes associated with cell development, differentiation, proliferation, survival, and angiogenesis [1]. STAT3 activation is detected in several malignancies, and its inhibition led to reversal of the malignant phenotype.

STAT3 activation has been described in nearly 70% of solid and hematological tumors [1]. STAT3 is constitutively activated by phosphorylation of Tyr705, in primary human pancreatic ductal adenocarcinomas (PDA), in PDA cell lines, and in PDA xenografts [2]. Studies in STAT3 conditional knockout mice demonstrate that the STAT3 pathway is inactive in normal pancreas, and it is not required for pancreatic development and homeostasis [2]. However, STAT3 was found to be necessary for the development of the acinar-to-ductal metaplasia process, an early event in PDA pathogenesis, which is mediated by ectopic expression of the Pdx1 transcription factor, a key regulator in early pancreatic cancer development [3]. Functional inactivation of STAT3 in a subset of PDA cell lines led to significant inhibition in cell proliferation *in vitro* and reduced tumor growth *in vivo*. Inhibition of activated STAT3 resulted in the delay of G1/S-phase progression due to inhibition of cyclin-dependent kinase 2 activity because of increased expression of p21/WAF1. Overall, the study clearly showed that with malignant transformation, activated STAT3 promotes proliferation of cells by modulating G1/S-phase progression and supports the malignant phenotype of human PDA cells [4].

Interestingly, activation of STAT3 also suppresses tumor growth [5] and induces differentiation and apoptosis in some contexts [6, 7]. The mechanisms underlying STAT3’s diverse and sometimes opposing roles are still largely unknown. It is assumed that STAT3 recruits specific co-activators and activates distinct gene expression programs based on the genetic background, type, and developmental stage of the cell [8]. This hypothesis raises an interesting issue of whether STAT3 pY705 and pS727 play a role in this process, considering their significance in STAT3-mediated control of gene transcription. Phosphorylation of Y705 is believed to be the key event in the transcriptional activation of STAT3 [9].

One of the transcriptional targets of STAT3 that is important in oncogenesis is MUC1 [10]. The promoter of MUC1 gene contains a STAT-responsive element, and STAT3 constitutively binds to it [11, 12]. Inhibition of STAT3 reduced the expression of MUC1 transcription and inhibited cellular motility in breast cancer cells [12]. The MUC1 glycoprotein that is aberrantly overexpressed in several epithelial cancers and is associated with transformation, tumorigenicity, invasion, and metastasis of carcinomas is an important downstream target of STAT3 [11, 13, 14]. Furthermore, MUC1-CT binds directly to JAK1 and STAT3 and promotes JAK1-mediated phosphorylation of STAT3. In turn, activated STAT3 induces expression of the MUC1 gene, in an auto-inductive loop. Therefore, it has been proposed that targeting STAT3 and MUC1 together may be a strategy for enhanced anti-tumor efficacy [15].

Since STAT3-MUC1 signaling is constitutively active in high-MUC1 tumor cells, we hypothesized that high-MUC1 cells are more sensitive to STAT3-inhibition. Although STAT3 has been a target for developing cancer therapy for a while, STAT3 inhibitors have not been successful in the clinic. For example, the STAT3 inhibitor Napabucasin (BBI608) was tested under phase III clinical trials for gastrointestinal (GI) cancers, including pancreatic cancer. However, the trial was discontinued due to futility [16]. To better understand the causes of treatment failure, the complex mechanisms of STAT3 signaling need to be elucidated. In this study, we aimed to understand the role of MUC1 in regulating differential phosphorylation of STAT3 and whether inhibiting STAT3 by Napabucasin interrupts MUC1-STAT3 interaction and downstream oncogenesis.

## Results

### STAT3 is overexpressed and correlates with poor overall survival in epithelial cancers

To assess the translational significance of STAT3-MUC1 interactions in epithelial cancers, we analyzed tumor data from The Cancer Genome Atlas (TCGA) and found that STAT3 was significantly overexpressed in majority of epithelial cancers vs normal (Figure 1A) and correlated with worse survival outcomes (Figure 1B). In addition, we found that STAT3 and MUC1 co-expression significantly correlated with reduced survival in gastrointestinal cancers (Figure 1C), thus showing the amplifying oncogenic effect that MUC1 and STAT3 have together.

**Figure 1.**
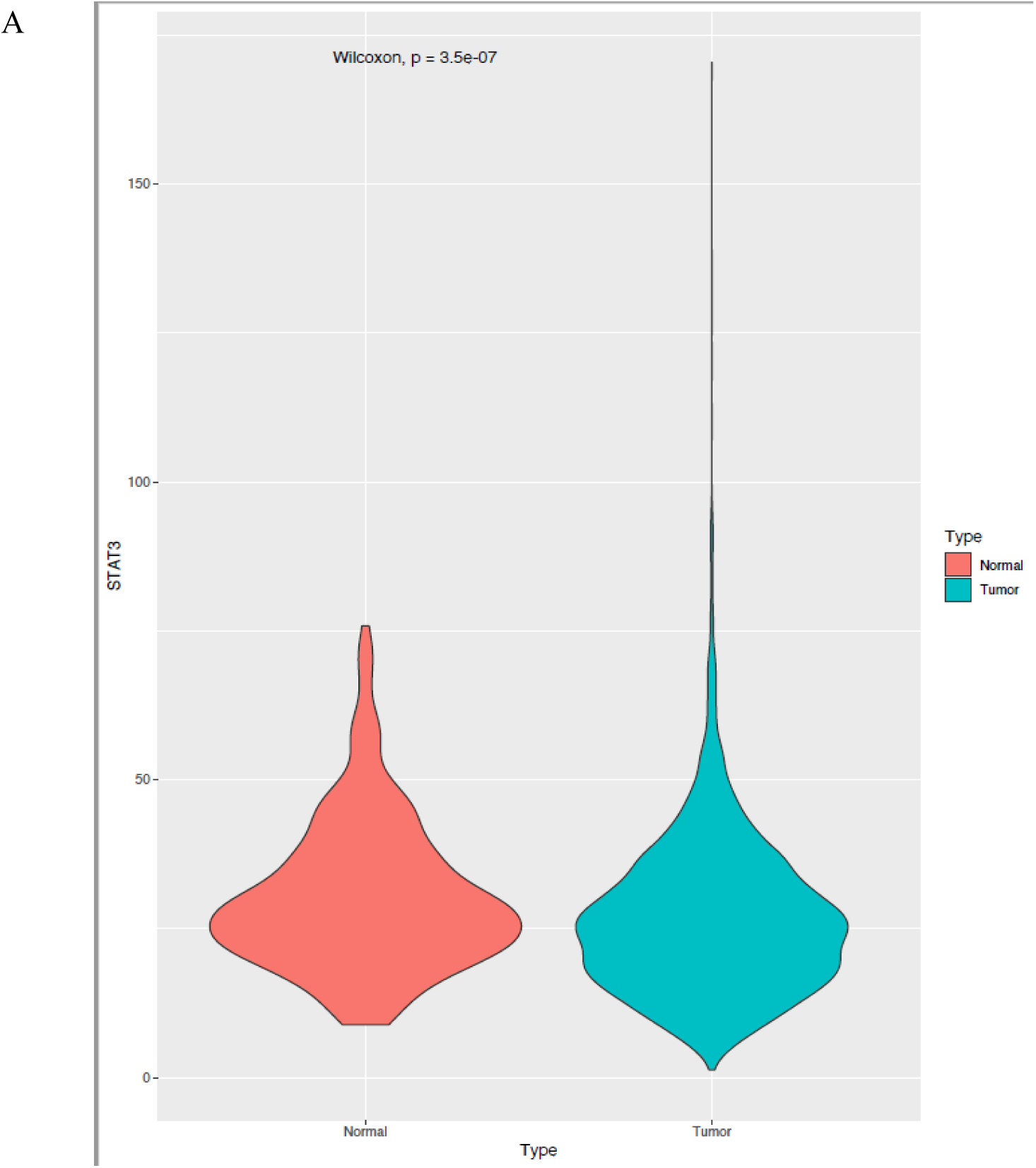
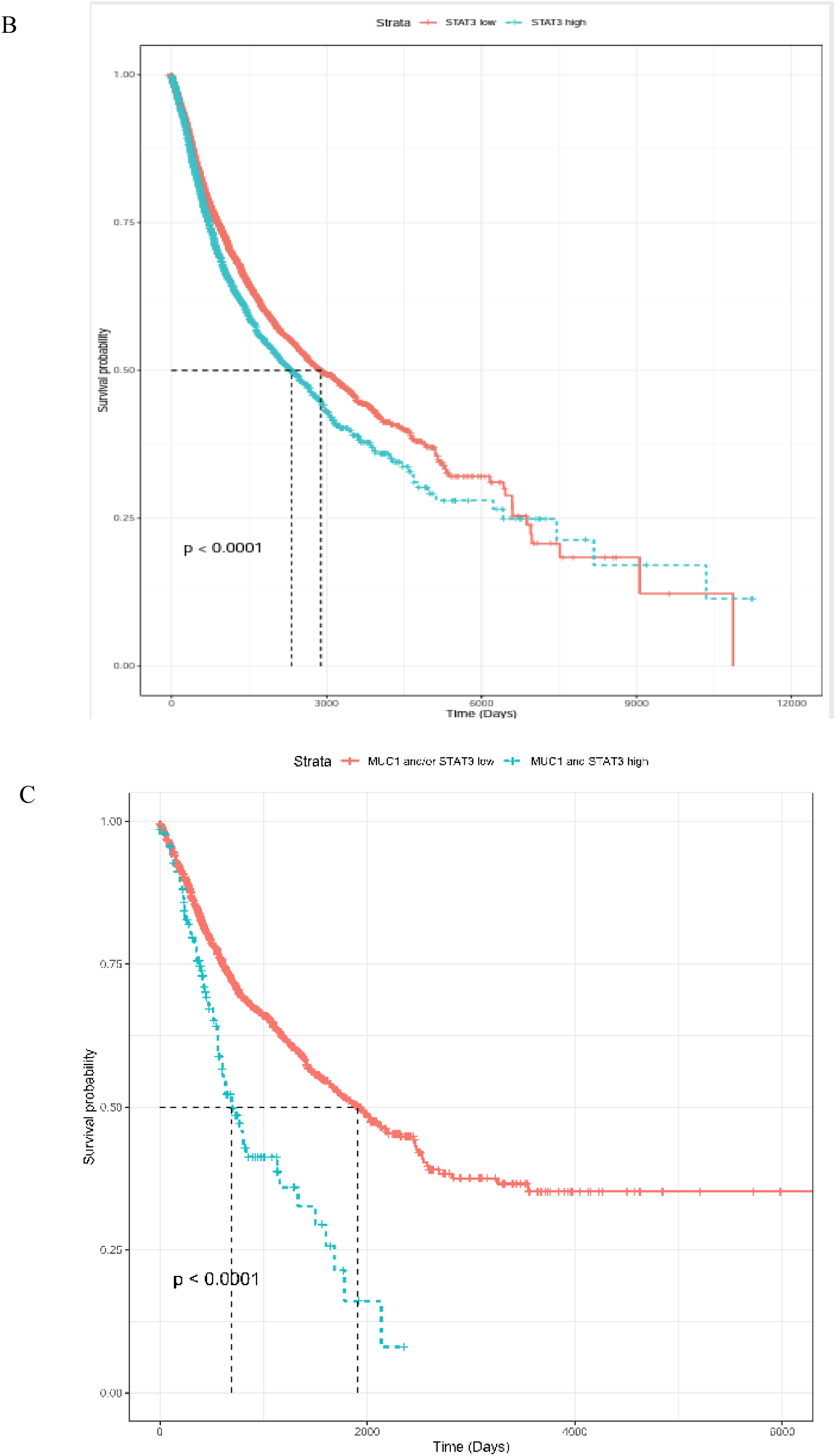
MUC1-STAT3 co-expression in epithelial cancers. **A.** Graph showing increased expression of STAT3 in epithelial tumor tissues (green) compared to normal (red). **B.** Kaplan-Meier survival plot showing correlation of STAT3 with overall poor prognosis in epithelial cancers (red-low STAT3; green-high STAT3). **C.** Kaplan-Meier survival plot showing correlation of MUC1 and STAT3 co-expression in gastrointestinal cancers (red-low MUC1 and STAT3; green-high MUC1 and STAT3).

### High MUC1 expressing tumor cells are more sensitive to the anti-tumor effects of Napabucasin compared to low MUC1 expressing cells

Since MUC1 and STAT3 co-expression led to worse outcomes, we hypothesized that MUC1 activates STAT3 through an unknown mechanism, amplifying its oncogenic effect. Therefore, we first aimed to study the pharmacodynamics of a STAT3-inhibitor, Napabucasin in low vs high MUC1 tumor cells *in vitro*. We used two human PDA cells lines that constitutively express high MUC1 (CFPAC and HPAFII) along with six pairs of isogenic human and murine tumor cell lines and stably transfected them with either an empty vector or full length MUC1, designated as “Neo” and “MUC1”, respectively. These isogenic cells included human pancreatic cancer cells MiaPaca2 and BxPc3, murine colon cancer cell line MC38, breast cancer cell line C57MG and ovarian cancer cell line MOVCAR. In addition, we used murine pancreatic cancer cell lines KCM (that express full-length human MUC1) and KCKO (genetically null for MUC1) [17, 18].

To determine whether MUC1 plays a role in STAT3 sensitivity to Napabucasin, we treated the panel of low and high MUC1 expressing cancer cells with increasing concentrations of Napabucasin. We found that the IC_50_ values of Napabucasin in PDA cells lines CFPAC and HPAFII (both expressing constitutively high MUC1) were similar (Figure 1A). However, in the subsequent experiments that used isogenic cell lines, we found that the IC_50_ values of the drug were significantly lower in MUC1 high cells (BxPC3.MUC1 and MC38.MUC1) than that in their low/no MUC1 expressing counterparts (BxPC3.Neo and MC38.Neo) (Figure 2B-left panel). Interestingly, when these cells were pretreated for 2 hours with 10µM of GO-203, a peptide blocking MUC1 homodimerization and therefore MUC1 signaling, we noted a reversal in the response to the drug (Figure 2B right panel), suggesting that MUC1 oncogenic signaling may be necessary for the drug to function effectively. This was further confirmed in the same cell lines using a single dose (0.8µM) of Napabucasin with and without pretreatment with GO-203 (Figure 2C). To further test our hypothesis that high MUC1 cells are more susceptible to the anti-tumor effects of Napabucasin, we tested the effect of the drug on a few more isogenic cells lines that express high and low/no MUC1 (MiaPaca-2 MUC1 and Neo; C57.mg.MUC1 and Neo; Movcar.MUC1 and Neo; and KCM and KCKO) (Figure 2D). All MTT data shown are at 48 hours post Napabucasin treatment. Taken together, the data convincingly showed that high expression of MUC1 in multiple tumor types sensitizes the cells to the anti-tumor effects of a STAT3 inhibitor. Furthermore, homodimerization of MUC1 CT may be necessary for the response to the drug. Thus, the data are in line with previous findings that MUC1 CT enhances STAT3 activation in cancer cells.

**Figure 2.**
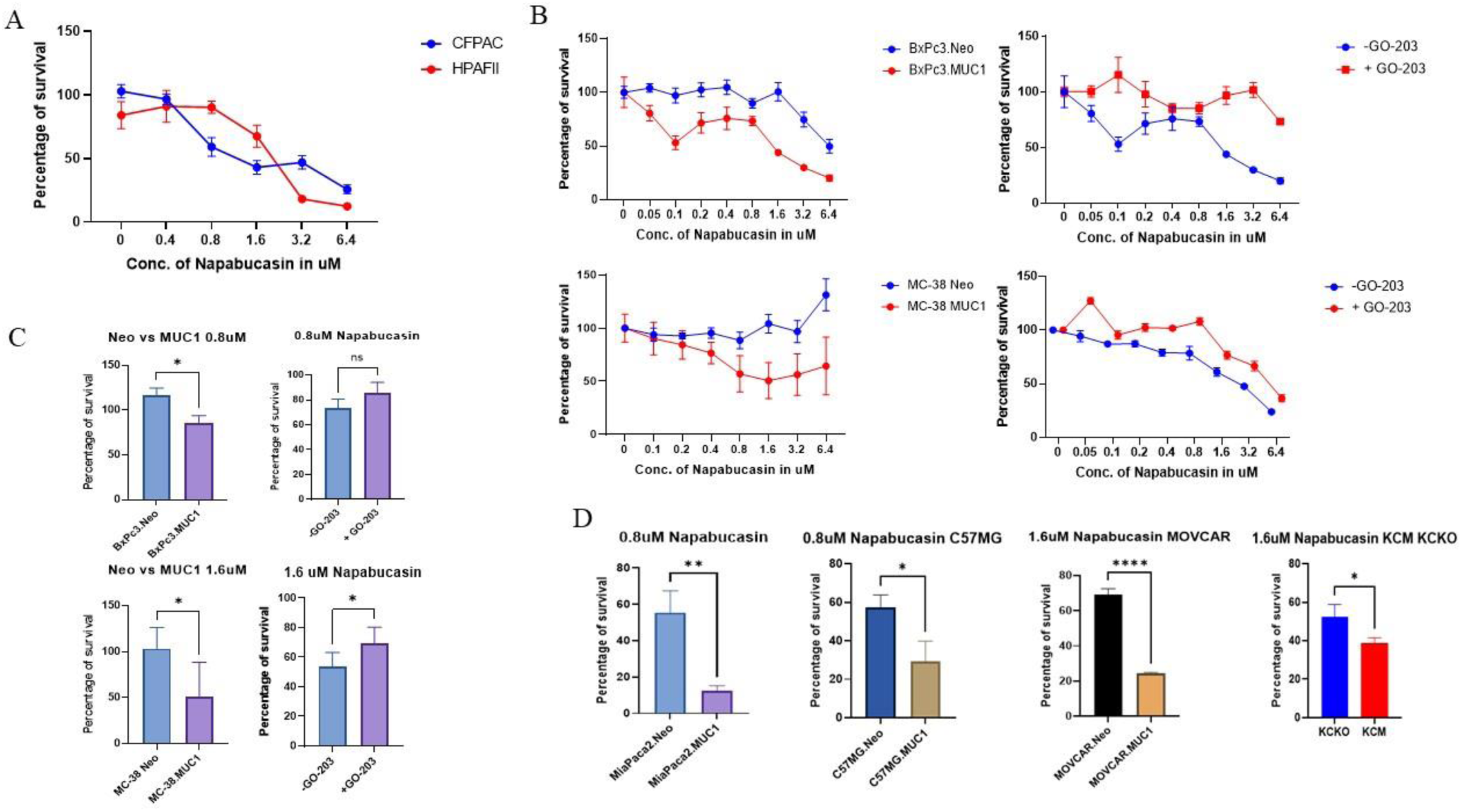
Napabucasin is more potent in high-MUC1 cells. **A.** Survival assay showing the kill curve of Napabucasin on human PDA cell lines CFPAC and HPAFII after 48 hours. **B.** Survival assay showing the kill curve on murine cancer cells transfected with either empty vector (Neo) or full-length MUC1 with or without 1-2 hours of pre-treatment with 10μM of MUC1-homodimerization blocking peptide GO-203 followed by the indicated concentrations of Napabucasin treatment for 72 hours have been shown. (Left) BxPc3.Neo and BxPc3.MUC1, BxPc3.MUC1 before and after treatment with GO-203 (top right), MC38.Neo and MC38.MUC1 (bottom left) and MC38.MUC1 before and after GO-203 treatment (bottom right) to increasing concentrations of Napabucasin. **C.** Survival assay showing the sensitivity of (top) BxPc3.Neo and BxPc3.MUC1 and (bottom) MC38.Neo and MC38.MUC1 to 0.8μM and 1.6 μM Napabucasin respectively after 72 hours, and that of BxPc3.MUC1 and MC38.MUC1 before and after treatment with 10μM of GO-203. **D.** Survival assay showing the sensitivity of MiaPaca2.Neo and MiaPaca2.MUC1, C57MG.Neo and C57MG.MUC1, MOVCAR.Neo and MOVCAR.MUC1 and KCKO and KCM cells to 0.8μM and 1.6 μM Napabucasin after 72 hours.

### MUC1 levels determine the phosphorylation status of STAT3

Phosphorylation of Y705 (pY705) is believed to be essential for STAT3 transcriptional activity. Previous studies have shown that phospho-S727 regulates STAT3 activity by enhancing dephosphorylation of Y705, triggering a multistep inactivation of STAT3 by rapid dissociation of pY705-SH2 through C-terminal tail modulation [19, 20]. To assess if MUC1 expression levels have any effect on differential phosphorylation of STAT3, we overexpressed full-length MUC1 in MiaPaca2 cells and knocked MUC1 down in HPAF II cells with a specific siRNA. We found that overexpression of MUC1 in MiaPaca2 cells slightly increased pY705 and decreased pS727 while knock down of MUC1 (MUC1 KD) in HPAFII cells decreased pY705 and slightly increased pS727 (Figure 3). Western blot and densitometry analysis is shown in Figure 3. Data suggest that MUC1 expression has some effect on the phosphorylation status of the Y705 and S727 and therefore plays a role in the subsequent activity of STAT3.

**Figure 3.**
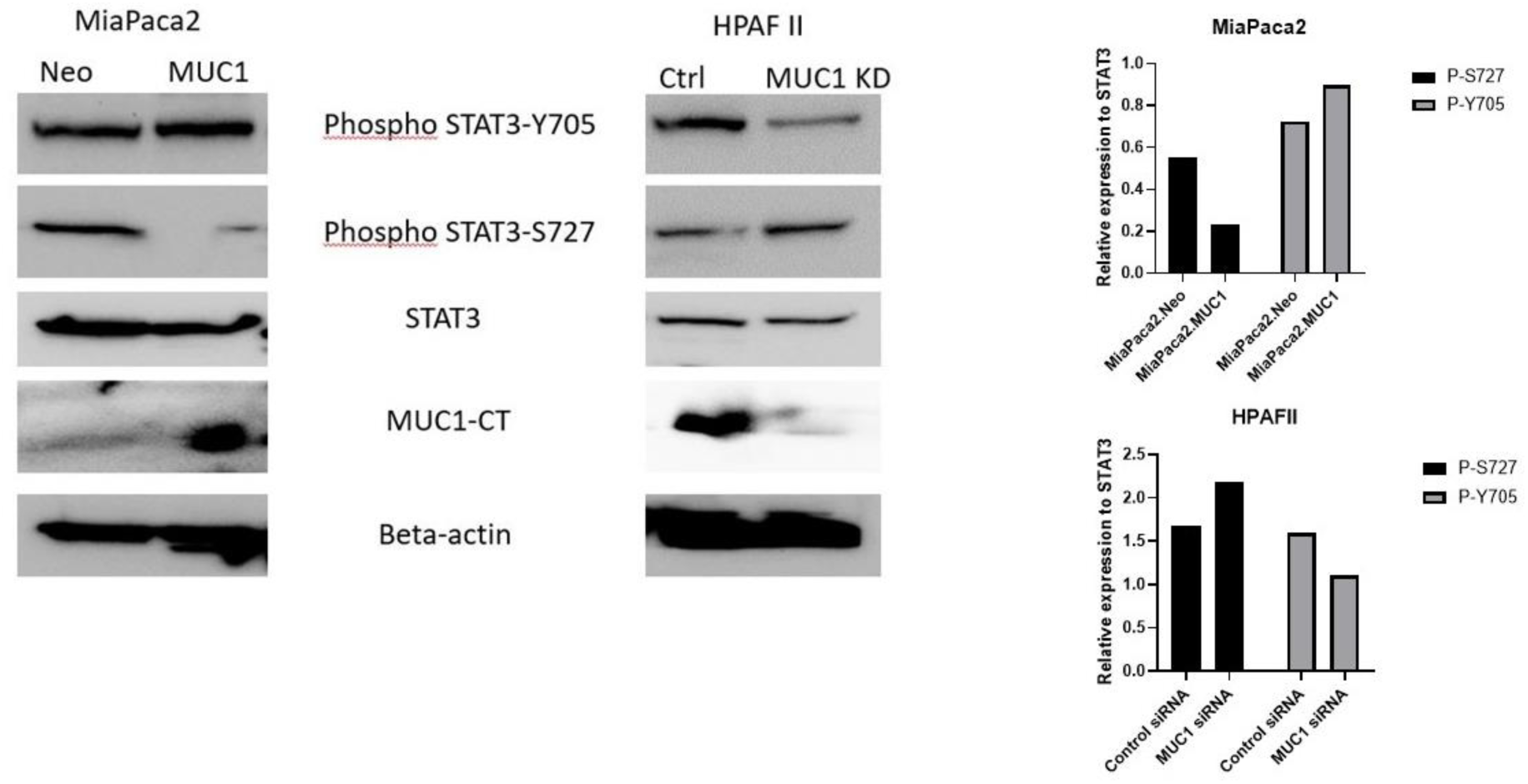
MUC1 levels determine the phosphorylation status of STAT3 in PDA cells. **A.** Western blot showing the expression of P-STAT3 Tyrosine 705 and Serine 727, total STAT3, MUC1-CT, and β-actin on (left) MiaPaca2.Neo and MiaPaca2.MUC1 cells and (right) HPAFII with control siRNA and HPAFII with MUC1 knockdown. **B.** Densitometric analysis of phosphorylated STAT3 S727 and Y705 in MiaPaca2.Neo and MiaPaca2.MUC1 cells (upper panel) and HPAFII.control siRNA treated and HPAFII MUC1.siRNA treated cells (lower panel).

### Napabucasin reduces phosphorylation of STAT3 at Y705 and disrupts STAT3-MUC1 interaction

To assess if Napabucasin has any effect on phosphorylation of STAT3 Y705 motif, and if that is affected by MUC1 expression level, we treated high and low MUC1 PDA cells (CFPAC and MiaPaca2 respectively) with 2µM of Napabucasin for 48 hours and measured the differences in phosphorylation of STAT3 at Y705, total STAT3 and MUC1 expression levels, and STAT3-MUC1 binding. After treatment with Napabucasin, high-MUC1 CFPAC cells had significantly lower levels of pSTAT3 at Y705, decreased STAT3 and MUC1 expression (Figure 4A), and reduced binding of STAT3 to MUC1 (Figure 4B).

**Figure 4.**
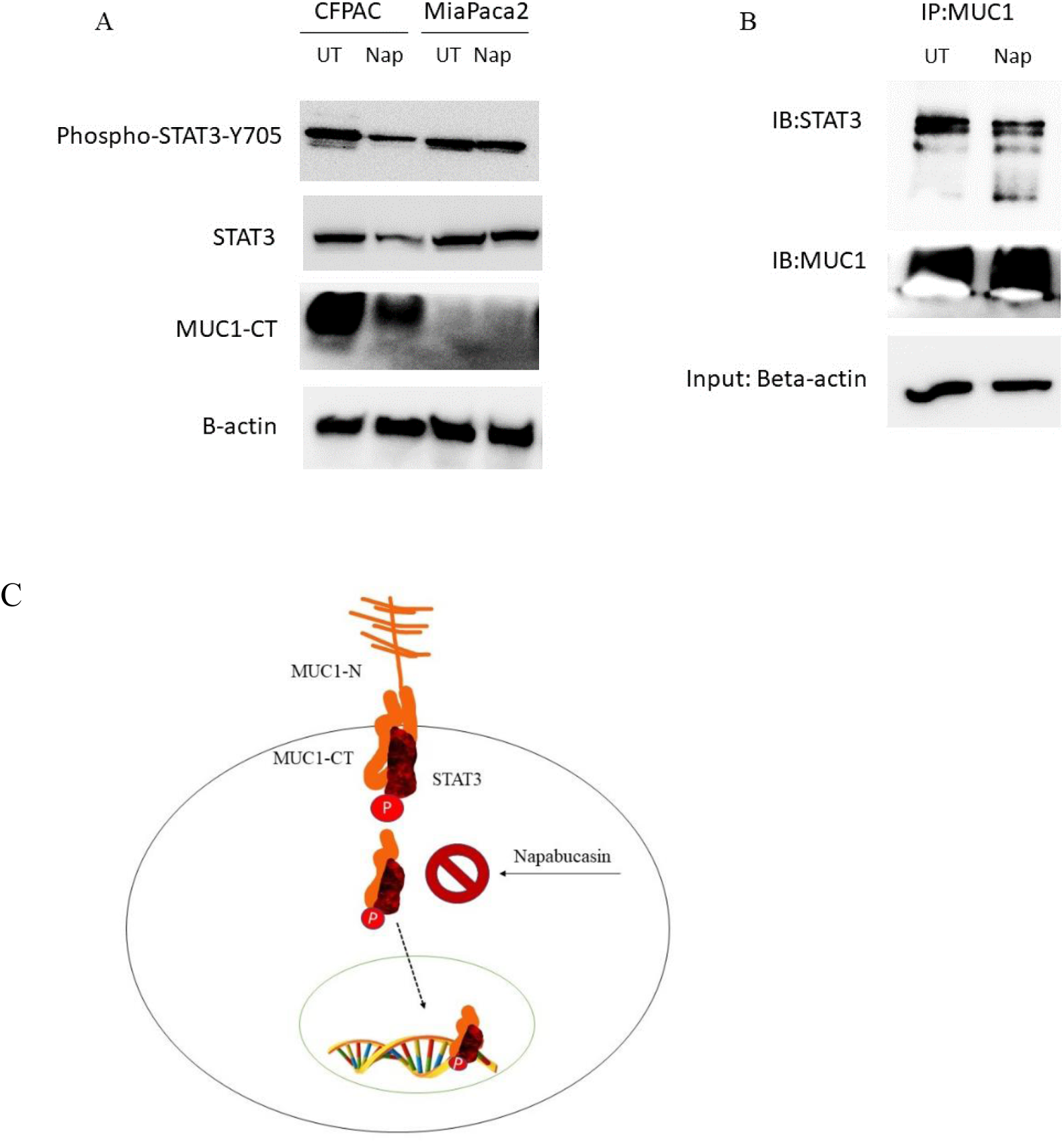
Napabucasin reduces STAT3 activity in high-MUC1 PDA cells. **A.** Western blot showing expression of P-STAT3 Tyrosine 705, STAT3, MUC1-CT in CFPAC and MiaPaca2 cells before and after treatment with 2 μM of Napabucasin for 48 hours. β-actin was used as endogenous control. **B.** Western blot showing expression of STAT3 and MUC1 in CoIP samples from CFPAC cell lysates before and treatment with 2 μM of Napabucasin for 48 hours. β-actin is shown as input. **C.** Proposed mechanism-of-action of Napabucasin in high-MUC1 cancer cells. Napabucasin inhibits the binding of MUC1 and STAT3, in turn, inhibiting their molecular crosstalk in high-MUC1 cells leading to cell death.

In contrast, we did not observe these effects in phosphorylation in the low MUC1 MiaPaca2 post Napabucasin treatment (Figure 4A). This supports our hypothesis that in high-MUC1 PDA cells, STAT3-MUC1 signaling is constitutively activated as a survival pathway and Napabucasin inhibits this pathway (Figure 4C).

### MUC1-STAT3 co-expression correlated to EMT and drug resistance genes in human tumor samples stratified into low and high MUC1 groups using the differential gene correlation analysis (DGCA)

To assess the clinical relevance of our data, we performed DGCA of STAT3 pathway genes with MUC1 gene expression. DGCA showed over 103 genes in the STAT3 pathway with differential correlation with MUC1 expression (Figure 5A). Genes that showed more than two-folds difference in correlation to MUC1 were mostly from the interleukin (IL) and interferon (IFN) families (Figure 5A). Pathway enrichment analysis of these genes showed that the top pathways altered in high vs low MUC1 samples include JAK-STAT pathway, EGFR inhibitor resistance and PI3K-Akt pathways amongst many others (Figure 5B). Since both MUC1 and STAT3 are known to regulate Epithelial-to-Mesenchymal Transition (EMT) and drug resistance, we were interested to find out the key genes that are involved in their partnership. We overlaid the Differentially Expressed Genes (DEGs) in MUC1 high vs low and MUC1-STAT3 dual high vs low with genes in the EMT and drug resistance pathways and found 11 DEGs in MUC1 high vs low samples illustrated in a Venn diagram (Figure 5C and D) and 13 DEGs in MUC1-STAT3 dual high vs low samples that are involved in both EMT, and drug resistance (Supplementary Figure 1). Since high MUC1 expression and high MUC1-STAT3 co-expression correlated to drug resistance, we wanted to focus on Napabucasin resistance specifically and find genes involved in this nexus. The molecule interaction network of Napabucasin was overlaid with the DEGs in low vs high MUC1 samples (Figure 5E) and MUC1-STAT3 dual low vs high samples from TCGA (Supplementary figure 2B). The molecular interaction network analysis of Napabucasin overlaid with DEGs in high vs low MUC1 showed predicted inhibition of all stemness markers that are upregulated in cancers (orange line with inhibition), namely STAT3, MUC1, ALDH1A1, Notch1, Yamanaka factors MYC, SOX2, KLF4 and more, and predicted activation of FAS, p38 MAPK, JNK, Caspase 3 and BAX that are usually downregulated (blue lines with arrowhead), out of which only SOX2 was significantly differentially expressed (Figure 5E). None of the DEGs from the PPI analysis in MUC1-STAT3 dual low vs high samples, were significantly differentially expressed in Napabucasin-molecular network. This could be due to lack of data on the molecules involved in Napabucasin signaling. Nevertheless, it emphasizes the importance of MUC1 in the Napabucasin network and highlights how it can influence potential outcomes to Napabucasin therapy. Our data corroborated with the predicted inhibition in this network showing that Napabucasin treatment reduced MUC1 levels in CFPAC cells (Figure 4A).

**Figure 5.**
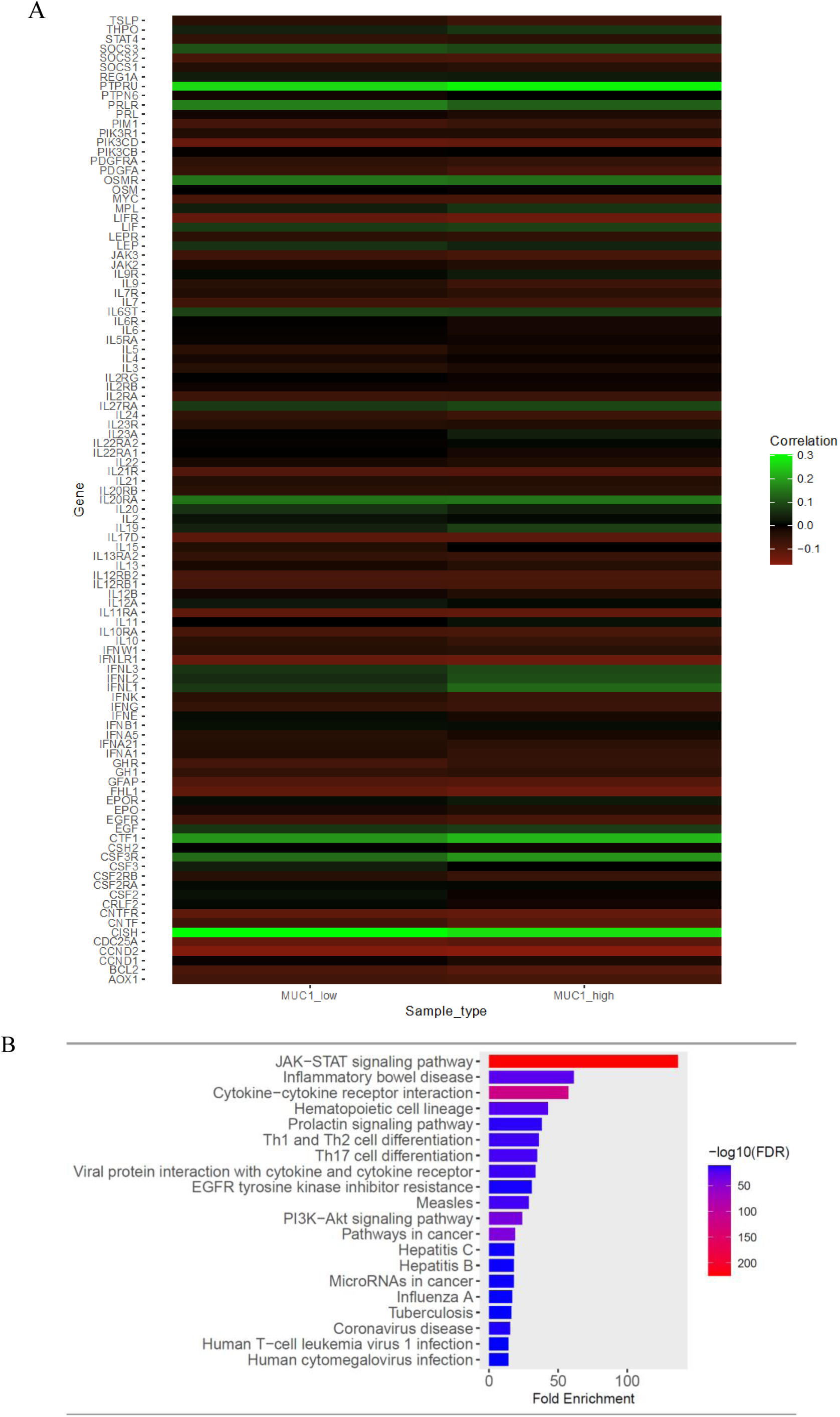
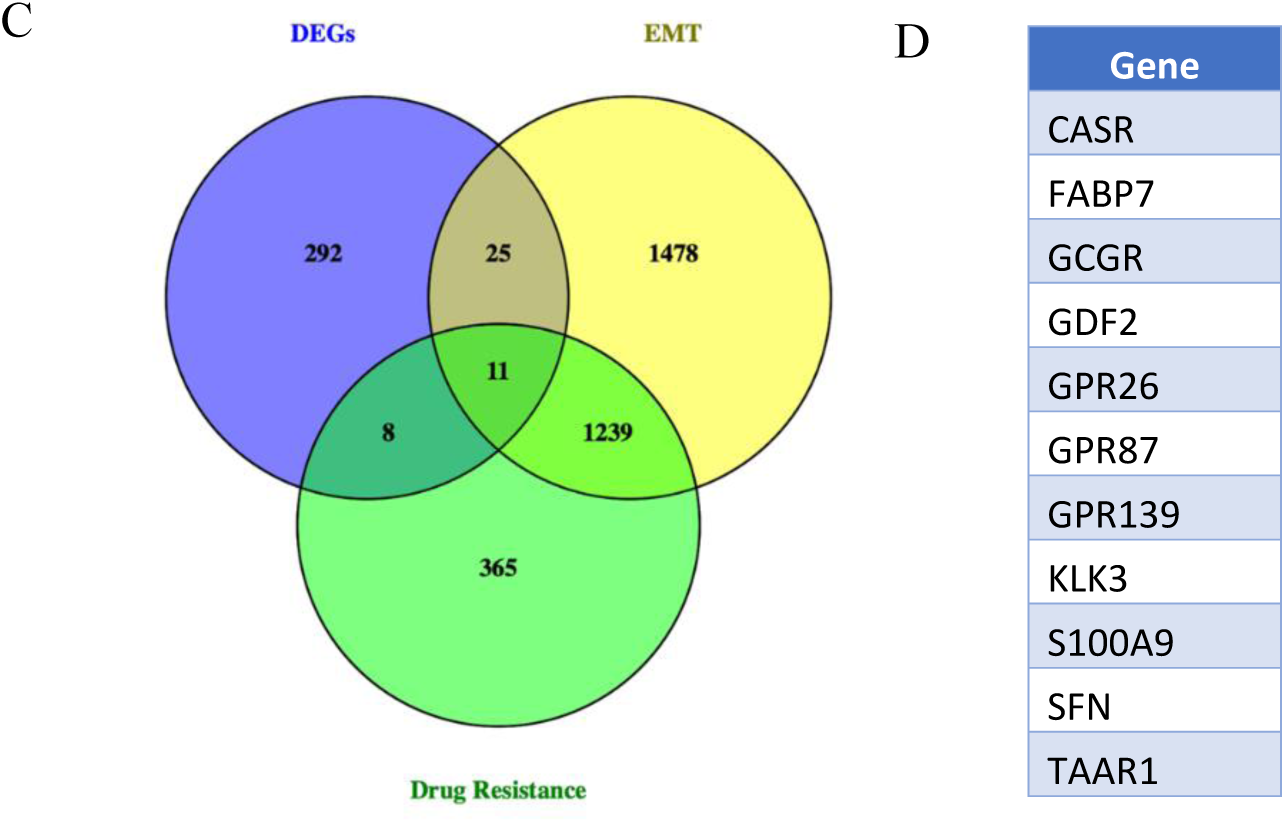
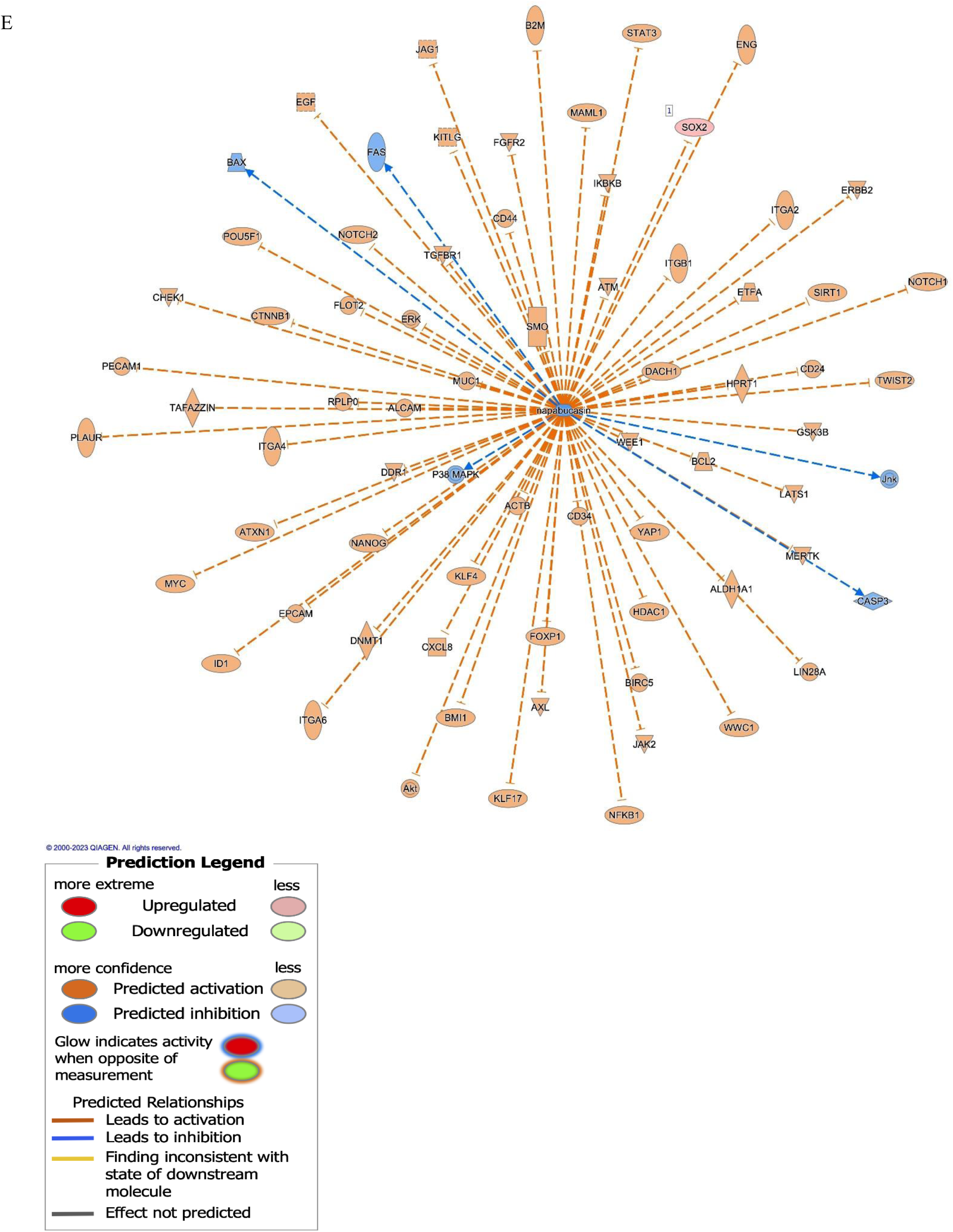
MUC1 and STAT3 co-expression is involved in EMT, drug-resistance and are part of the Napabucasin network. **A.** Heatmap showing gene correlation of STAT3 pathway genes and MUC1 gene expression. Differential gene correlation analysis results from MUC1 low vs. high gene expression in 13,509 TCGA tumor samples were used to visualize difference in correlation with STAT3 pathway genes (103 pathway genes remained in results from filtering/analysis).**B.** Pathway enrichment analysis with the differentially expressed genes from **A** using ShinyGO 0.76.3 **C.** The Venn diagram displays the common genes between DEGs from MUC1 low vs. high in BRCA, PAAD, CESC, LIHC, and OV TCGA samples with EMT and drug resistance related pathways. **D.** List of genes common between EMT and drug resistance. **E.** Molecule interaction network for Napabucasin overlaid with DEG results from MUC1 low vs. high in BRCA, PAAD, CESC, LIHC, and OV TCGA samples. The Napabucasin molecule interaction network displays the relationship between the drug and targeted molecules. The DEG results from MUC1 low vs. high in BRCA, PAAD, CESC, LIHC, and OV TCGA samples were overlaid on to the network to display predicted impact. In this dataset, SOX2 is observed to be significantly differentially expressed (shown in red). Line colors represent the predicted activation state of the molecule (orange represents predicted activation and blue is predicted inhibition).

## Discussion

STAT3 has been a long-standing target for developing cancer therapy, including GI cancers. Although current standard treatments for GI cancers are somewhat efficient, recurrence is still inevitable, especially in PDA [21]. Many studies have indicated that presence of pancreatic cancer stem cells (PSCs) may be a major cause of disease relapse [22]. Therefore, targeting PSCs might be a promising strategy to eradicate PDA. Given the well-documented role of overactive STAT3 signaling in maintaining stemness in PDA, the therapeutic potential of targeting this pathway should be emphasized. Multiple attempts to develop inhibitors against STAT3 pathway have been reported, and a variety of STAT3 inhibitors, including chemicals, STAT3-binding peptides, and siRNA reagents, have been developed with various degrees of success [23, 24].

Napabucasin (BBI608) is a novel STAT3-specific inhibitor identified by Li *et. al.* [25]. The authors revealed that Napabucasin efficiently suppressed metastasis and relapse of a variety of cancers by inhibition of STAT3-driven gene transcription. Importantly, Napabucasin treatment impaired spheroid formation of liver cancer stem cells and downregulated the expression of stemness genes such as SOX2, BMI-1, Nanog, and c-Myc. Considering the promising preclinical data of Napabucasin as both a monotherapy and in combination with conventional chemotherapeutic methods, several clinical trials have been performed [26]. Furthermore, a phase III trial of Napabucasin for refractory colorectal cancer highlighted STAT3 as an essential target for the treatment of patients with elevated pSTAT3 expression [27]. However, to improve outcomes in the clinic, it is crucial to find subpopulations of tumors that will be more sensitive to Napabucasin. In 2009, MUC1 was ranked as the second most targetable antigen to develop cancer vaccines by the National Cancer Institute [28]. Since then, a plethora of studies has shown the oncogenic role of MUC1 in increasing stemness, modulating drug resistance, as well as regulating signaling pathways in epithelial cancers [14, 29]. MUC1 is known to correlate with poor prognosis in pancreatic cancer (https://www.proteinatlas.org/ENSG00000185499-MUC1/pathology). Since MUC1 and STAT3 were found to regulate each other’s expression in an auto-inductive loop, we hypothesized that in high-MUC1 cancer cells, the STAT3-MUC1 pathway is constitutively activated as a survival pathway and therefore, high-MUC1 cells will be more sensitive to the anti-proliferative effect of STAT3-inhibitor Napabucasin. To assess the clinical relevance of our hypothesis, we analyzed RNA sequencing data across some epithelial cancers from TCGA and found that STAT3 is overexpressed and correlates with overall poorer survival in these cancer types (Figure 1 A and B). In addition, we found that co-expression of MUC1 and STAT3 correlated to even poorer overall survival in gastrointestinal cancers, indicating that MUC1 enhances the oncogenic activation of STAT3.

We confirmed that Napabucasin significantly inhibited survival of the tumor cells with high MUC1 expression compared to low MUC1 cell lines (Figure 2). We found that MUC1 overexpression results in increased phosphorylation of STAT3 at its activation site Y705 with a concomitant decrease in phosphorylation at its degradation site S727. Importantly, when MUC1 is knocked down, the reverse is observed (Figure 3). These data indicate that MUC1 stabilizes STAT3 by reducing its degradation signal P-S727 (Figure 3).

Following treatment with Napabucasin, the phosphorylation of STAT3 at Y705 and total STAT3 levels were decreased in high-MUC1 CFPAC cells but not in low-MUC1 MiaPaca2 cells (Figure 4A). Napabucasin treatment reduced binding of MUC1 to STAT3 in high-MUC1 CFPAC cells (Figure 4B). This indicates that in high-MUC1 cells, STAT3 is stabilized by MUC1 with increased phosphorylation at Y705, that maintains the STAT3-MUC1 auto-inductive loop. One study [30] found the underlying mechanism of downregulated STAT3 protein levels was mediated by protein synthesis inhibition induced by Napabucasin. Here, we find that Napabucasin significantly impairs the binding of STAT3 and MUC1, thus breaking the auto-inductive loop which is a novel pharmacological mechanism-of-action of Napabucasin.

We also identified more than 100 DEGs in the STAT3 pathway in high vs low MUC1 tumors (Figure 5C). These DEGs in high vs low MUC1 tumors were associated with EMT and drug resistance (Figure 6A). PPI analysis of these DEGs when overlaid with Napabucasin molecular interaction network revealed that SOX2 was significantly differentially expressed. The role of SOX2 in the Napabucasin network needs to be functionally elucidated in the future. Another limitation of this study is the lack of *in vivo* data showing that high-MUC1 cancer cells respond better to Napabucasin treatment compared to isogenic low MUC1 cells. However, analysis of clinical data from TCGA corroborates with our hypothesis that MUC1 and STAT3 co-expression correlate to increased drug resistance and poor survival. A recent study reported that Napabucasin treatment inhibited the STAT3-MUC1 pathway in stemness-high cells and that high MUC1 status in these cells was associated with Paclitaxel resistance. It provided a new mechanism for the association between cancer stemness and drug resistance. This study also showed how combining Paclitaxel with Napabucasin could be a promising strategy to combat cancer [31]. To our knowledge, this is the first study to evaluate the role of MUC1 in differential phosphorylation and regulation of STAT3 in the context of Napabucasin’s efficacy. In our study, we reveal that Napabucasin treatment reduced activation of STAT3 and MUC1 as well as their binding, and therefore is more potent against high-MUC1 cells. Overall, our results support the potential use of Napabucasin as an efficacious anti-tumor therapeutic agent with a probability of better outcome in high-MUC1 tumor sub-populations, singly or in combination with other therapeutic agents.

**Figure 6.**
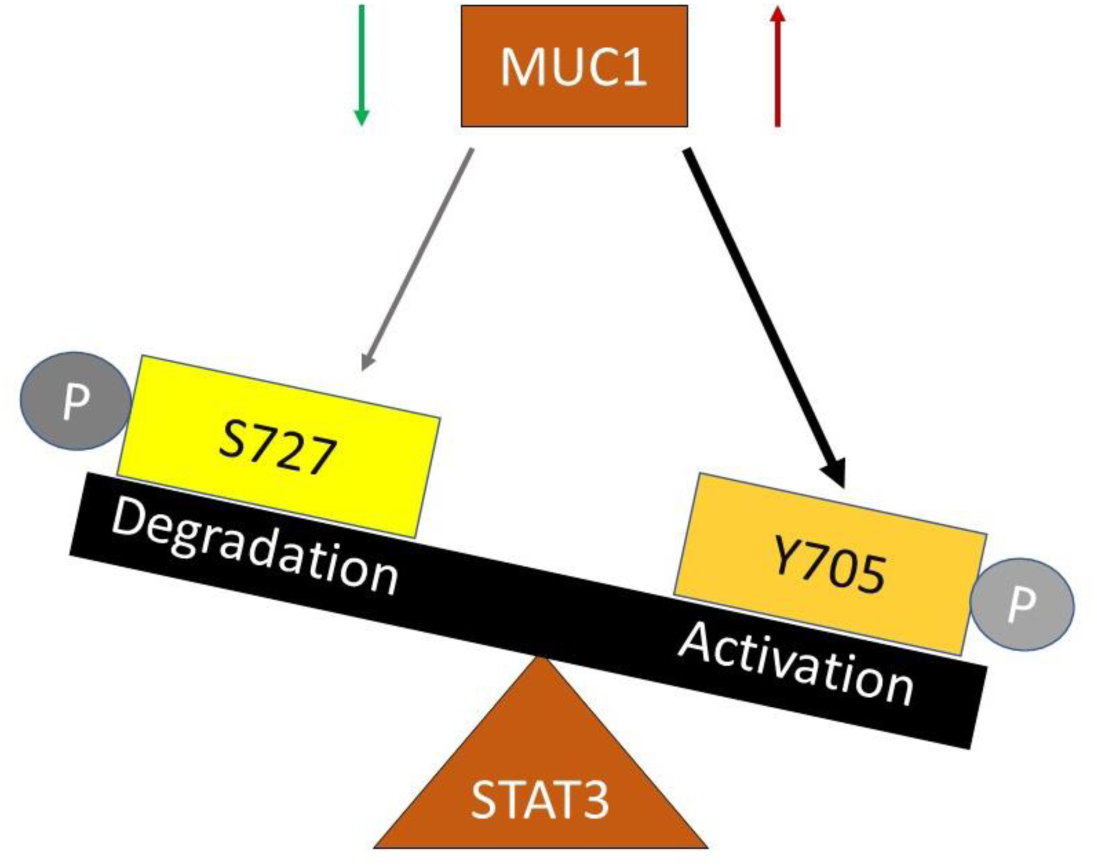
Proposed mechanism of regulation of STAT3 activity by MUC1. Overexpression of MUC1 induces phosphorylation of STAT3 at the Tyrosine 705 residue that is associated with increased activity and MUC1 downregulation induces phosphorylation of STAT3 at the Serine 727 residue that is associated with degradation.

## Materials and Methods

### TCGA Data Analysis

#### TCGA Gene Expression Analysis

RNA-sequencing data from 7,572 epithelial cancer samples available from TCGA was analyzed and STAT3 expression values in normal and tumor samples were plotted. The types of cancers include breast, lung, endometrium, kidney, head-neck, thyroid, prostate, colon, stomach, bladder, liver, cervix, pancreas, esophagus, adrenal, and gallbladder cancer. Plot was created using ggplot2 (version 3.3.5) package in R (version 3.6.3).

#### Gene Correlation Analysis

Tumor samples from TCGA cancer projects were split into MUC1 low/high group based on the MUC1 gene expression relative to the average MUC1 gene expression in normal samples. Datasets was filtered to remove lowest expressed genes by the dispersion measure. Differential gene correlation analysis was performed on these two groups using DGCA (Differential Gene Correlation Analysis) in R (version 3.6.3). Gene correlation analysis was run, using the Benjamini-Hochberg p-adjustment measure. STAT3 pathway genes were selected and their correlation values with MUC1 were visualized in a heatmap, using ggplot2 (version 3.3.5) package in R.

#### Molecular Interaction Network Analysis

The drug network figure represents Napabucasin, and associated molecules overlaid with IPA predictions on activation states. The Molecule Activity Predictor feature in IPA overlaid DEG results from MUC1 low vs. high in BRCA, PAAD, CESC, LIHC, and OV TCGA samples on to the network to display predicted impact. The SOX2 gene, in red to show increased expression, is a DEG that is associated with Napabucasin. Other molecules are colored orange for predicted activation or blue for prediction inhibition activity. The relationship arrows were generated to match the predicted activation states.

#### Survival Analysis

Survival analysis for STAT3 and overall survival in 13,509 cancer samples were computed using the Kaplan-Meier estimate and plots were made using ggplot2 (3.3.5) package in R (3.6.3). The average expression for STAT3 in normal samples was used to determine if tumor samples were low or high in STAT3 expression.

Survival analyses for MUC1/STAT3 expression and overall survival in 2,055 gastric cancer type samples were computed using the Kaplan-Meier estimate and plots were made using ggplot2 (3.3.5) package in R (3.6.3). The average expression for MUC1/STAT3 in normal samples was used to determine if tumor samples were low or high in MUC1/STAT3 expression.

### Cell lines and culture

Human PDA cell lines (CFPAC, HPAF-II and MiaPaca2) and murine cancer cell lines MC38 (colon), C57MG (breast), MOVCAR (ovarian) were obtained from American Type Culture Collection and KCM and KCKO (pancreatic) cells were generated as described [17, 18] and cultured as instructed. Cell lines were maintained in Dulbecco’s Modified Eagle Medium (DMEM; Gibco). All media was supplemented with 10% fetal bovine serum (FBS; Gibco or Hyclone), 3.4 mM l-glutamine, 90 units (U) per ml penicillin, 90 μg/ml streptomycin, and 1% non-essential amino acids (Cellgro). Cells were kept in a 5% CO_2_ atmosphere at 37℃. MUC1 WT sequence was cloned into the pLNCX.1 vector consisting of the neomycin resistance gene (neo) and confirmed by DNA sequencing. Neo cells had the empty vector with the G418 resistance gene (neo) and MUC1 cells had the full length MUC1 gene and G418 resistance gene (neo). All cells with Neo and MUC1 were generated by transfection with Lipofectamine 3000 (Thermo Fisher) according to the manufacturer’s protocol and maintained in medium containing Geneticin (G418; Invitrogen, Carlsbad, CA, USA) [17]. Every passage of Neo or MUC1 transfected cells were maintained in a final concentration of 150 μg/ml of the antibiotic G418 (50 mg/ml) (Thermo Fisher) to ensure positive selection. HPAFII cells were serum-starved for 24 hours and then treated with control siRNA from Life Technologies or MUC1 siRNA from Perkin Horizon according to the respective manufacturer’s protocol using Lipofectamine RNAiMAX Transfection Reagent (Thermo Fisher Scientific) for 72 hours. For all experiments, cell lines were passaged no more than 10 times.

### Treatment With Napabucasin and Western Blotting

The cell lines used were MiaPaca2. Neo, MiaPaca2. MUC1, CFPAC, HPAFII. control siRNA, and HPAFII. MUC1siRNA. Cells were treated with either 2 μM Napabucasin (Selleckchem, USA) or the vehicle (phosphate buffer saline) for 48 hrs. Cell lysates were prepared and western blotting performed as previously described [29]. Membranes were blocked with commercial blocking buffer (Thermo Fisher) for 30 min at room temperature and incubated with primary antibodies overnight at 4°C. The antibodies used were: Armenian hamster monoclonal anti-human MUC1 cytoplasmic tail (CT2) antibody (1:500). MUC1 CT antibody CT2 was originally generated at Mayo Clinic and purchased from Neomarkers, Inc. (Portsmouth, NH) [32]. CT2 antibody recognizes the last 17 amino acids (SSLSYNTPAVAATSANL) of the cytoplasmic tail (CT) of human MUC1. Membranes were also probed with the following antibodies from Cell Signaling Technology (1:1,000), p-STAT3 (Y705), total STAT3, β-actin, and from ABclonal (1:1000) p-STAT3 (S727) and β-actin. Densitometric analysis was conducted using the ImageJ software. First, each density unit for the particular protein was normalized to their respective β-actin density and then represented as phospho/total.

### Co-Immunoprecipitation

PDA cells were serum-starved for 24 hours and treated with 2μM Napabucasin for 48 hours. After that, the media was removed, cells were washed with PBS and lysate was collected with complete lysis buffer (Lysis buffer with 1X Halt Protease and Phosphatase inhibitor), with the help of a cell scraper. The lysate was vortexed briefly and then sonicated and kept on ice for 10 minutes, followed by centrifugation at 14,000 RPM for 15 minutes at 4°C. The supernatant was collected in a fresh tube, BCA assay was performed by Pierce BCA Assay kit (Thermo Fisher Scientific) to estimate protein concentration as per the manufacturer’s protocol. For co-immunoprecipitation, 1 mg of lysate was used to pull down MUC1 with 100 μg of MUC1-CT2 antibody or an anti-Armenian Hamster IgG control antibody using the Pierce Co-IP kit (Cat. No. 26149). All the steps were performed according to the manufacturer’s protocol. For WB, the eluate was mixed with 5X SDS sample loading buffer supplied by the manufacturer and run on an SDS gel and protocol for WB was followed.

### MTT Assay and addition of MUC1 blocking peptide

5,000 cells were plated in 96 well plates and allowed to grow overnight. Next day, the cells were treated either with PBS or increasing concentrations of Napabucasin in triplicates for 24-96 hr. Then 20μl of MTT solution (5 mg/ml) was added to each well and incubated for 3-4 hours at 37℃. Following that, the media with MTT was removed and 200μl of DMSO was added to each well to dissolve the formazan crystals for 10 min and the O.D. was measured with a plate reader (Multiskan, Thermo Fisher) at 560 nm. For blocking MUC1 signaling, the peptide GO-203 was added to the cells 2 hours before treatment with Napabucasin. 48h data is shown.

### Statistical Analysis

Differences between groups were examined using unpaired two-tailed Student’s t-tests, one-way and two-way ANOVAs. Statistical comparisons were made using the GraphPad Prism 9.0. p-values of <0.05 were considered statistically significant (*p < 0.05; **p < 0.01; ***p < 0.001; ****p < 0.0001).

## Acknowledgements

We would like to thank Ms. Sophia Shwartz, Ms. Priyanka Lala, and Mr. Manuel Rodriguez Cardona for their help in the lab.

## Author contributions

M.B. conceived the idea, performed investigation, experiments, data analysis and prepared the manuscript; A.V. and A.H. performed experiments; A.S. performed all bioinformatics data analysis with inputs from M.B. and P.M.; P.M. provided scientific input, supervised the work, revised the manuscript, and provided funding; C.B. provided supervision and funding.

## Data availability statement

All of the RNA sequencing data analyzed was downloaded from the Genomics Data Commons Portal: https://portal.gdc.cancer.gov.

