## Supplementary information for "Overexpression of MUC1 influences the anti-proliferative effect of STAT3-inhibitor Napabucasin in epithelial cancers"

A

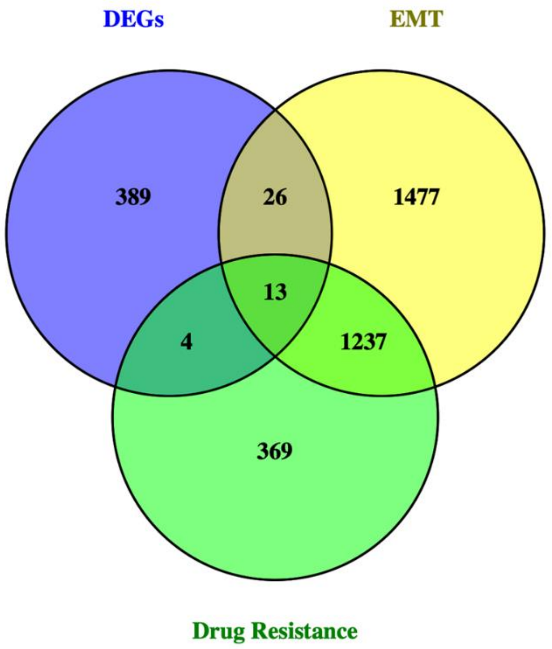


B

| Gene |
|---|
| BCYRN1 |
| CASR |
| CCKBR |
| FABP7 |
| FFAR1 |
| GPR12 |
| GPR50 |
| GPR139 |
| IGF2 |
| KLK1 |
| OPRK1 |
| PCDH10 |
| PRKCG |

**Supplementary Figure 1. A.** DEGs from MUC1/STAT3 low vs. MUC1/STAT3 high in BRCA, PAAD, CESC, LIHC, AND OV TCGA samples. The venn diagram represents the common genes involved with MUC1/STAT3 low vs. MUC1/STAT3 high DEGs and the EMT (epithelial-mesenchymal transition) and drug resistance pathway genes. Genes from multiple IPA pathways related to EMT and drug resistance were included in each group. **B.** Intersection of DEGs from MUC1/STAT3 low vs. MUC1/STAT3 high in BRCA, PAAD, CESC, LIHC, and OV TCGA samples and genes in the EMT and drug resistance pathways
